# Persistence of residual Anopheles gambiae populations associated with frontier human settlements and nocturnal livelihood-related activities at the fringes of a large conservation area in southern Tanzania

**DOI:** 10.1101/2025.10.24.684313

**Authors:** Deogratius R Kavishe, Lucia J Tarimo, Rogath V Msoffe, Katrina A Walsh, Lily M Duggan, Fidelma Butler, Emmanuel W. Kaindoa, Gerry F Killeen

## Abstract

Following the rapid scale-up of long-lasting insecticidal nets (LLINs) in 2008 across the Kilombero Valley of southern Tanzania, *Anopheles gambiae* Giles, a highly efficient and human-specialized malaria vector, essentially disappeared within two years and has rarely been detected since. However, an ecological study of its sibling species within the *An. gambiae* complex, namely *An. arabiensis* Patton and *An. quadriannulatus* Theobald, in an area spanning human settlements and conserved wilderness, nevertheless detected sparse, highly focal populations of *An. gambiae*. Out of 4,704 *An. gambiae* complex specimens collected, only 10 were identified as nominate *An. gambiae* by polymerase chain reaction. Seven of these were captured at three surveyed fishing camps inside a Wildlife Management Area (WMA), while none were caught in five established villages west of the WMA or the national park to the east. Two of the other three were found among scattered farming settlements along the edge of the WMA. The final individual was caught in a near-natural area deep inside the WMA, but even this site was near small transient homesteads. Poisson regression modelling confirmed that capture rates at fishing camps were higher than in the peripheral settlements (Relative Rate [RR] [95% Confidence Interval (CI)] = 0.088 [0.011, 0.695], P = 0.021) or conserved natural areas (RR [95% CI] = 0.043 [0.003, 0.495], P = 0.012). Conservative estimates put the mean standing populations at each of the three fishing camps between 26 and 41 females per settlement.

The absence of *An. gambiae* from well-conserved areas without resident humans aligns with its anthropophagic nature and dependency on human hosts. Residual *An. gambiae* populations persisting among frontier communities appear linked to locally common, livelihood-related nocturnal activities that preclude the practical use of LLINs, specifically fishing and guarding crops against wildlife. The continued presence of this previously important species in these ecological niches, where local livelihoods apparently undermine high LLIN coverage, shows persistence under sustained, partial insecticide pressure. This could lead to the re-emergence of this vector through the gradual evolution of resistance. However, their sparse and focal distribution suggests that new supplementary interventions, such as transfluthrin emanators, might effectively eliminate these remaining refuge populations.

## Introduction

Long-lasting insecticidal nets (LLINs) and indoor residual spraying (IRS) have proven highly effective interventions against malaria transmission. Their effectiveness stems from targeting insecticides to the most anthropophagic and efficient vector mosquitoes (Diptera: Culicidae) that prefer human blood and seek hosts sleeping indoors at night (Killeen, 2014; Killeen, Kiware, et al., 2017). In many contexts, LLINs and IRS have even achieved local elimination of some highly human-dependent African vectors, namely the nominate sibling species of the *Anopheles funestus* group and *An. gambiae* complex (Hosack et al., 2025; Killeen, Kiware, et al., 2017; Killeen et al., 2013; Sinka et al., 2016). Following the scale up of LLINs across Africa over 15 years ago, *An. gambiae* Giles *sensu stricto* rapidly disappeared from large tracts of its historical range (Hosack et al., 2025; Sinka et al., 2016), including some particularly well studied parts of Kenya (Bayoh et al., 2010; Mwangangi et al., 2013) and Tanzania (Russell et al., 2011).

One such historically well characterized malaria transmission system is the Kilombero Valley in southern Tanzania, the largest low-lying flood plain in east Africa (Charlwood et al., 1995; Killeen & Smith, 2007; Smith et al., 1993; Smith et al., 1995). Before bed nets were introduced, malaria transmission in the Kilombero Valley was intense, with ∼70% parasite prevalence among children and entomological inoculation rates (EIRs) of hundreds or even thousands of infectious bites per person per year recorded in the 1990s (Smith et al., 1993). This intense transmission was primarily driven by the highly anthropophagic and endophilic vector *Anopheles gambiae s.s.*, which exhibited such an apparently strong dependence upon human blood that it was less abundant than *An. arabiensis* Patton in relative terms wherever the latter partially zoophagic species could access cattle (Charlwood & Edoh, 1996; Charlwood et al., 1995; Killeen et al., 2001; Tirados et al., 2006; White, 1974; White et al., 1972). In the early 2000s, Tanzania rapidly scaled up vector control measures, starting with targeted distribution of insecticide-treated nets to young children and pregnant women (Abdulla, 2001; Killeen, Tami, et al., 2007; Mushi, 2003), but then progressing towards universal coverage of all age groups with the specific purpose of suppressing vector populations (Hawley et al., 2003; Killeen, Smith, et al., 2007) after this target was adopted as global policy in 2007 (WHO, 2007). In the Kilombero Valley, however, high usage rates across the entire population had already been established and sustained for several years, although the bed nets in question were largely untreated in practical terms (Abdulla, 2001; Killeen, Tami, et al., 2007). A large-scale effectiveness evaluation carried out across the valley in the late 1990s demonstrated that social marketing, combined with targeted partial subsidies for vulnerable population groups through discount vouchers, was highly effective as a means of promoting bed net delivery through private sector retailers (Abdulla et al., 2005; Marchant et al., 2002; Mushi, 2003; Nathan et al., 2004; Schellenberg et al., 2001; Schellenberg et al., 1999). Beyond the term of this operational research study, subsequent scale up of new delivery mechanisms at national level in the early 2000s successfully sustained high net coverage of all age groups across the valley (Killeen, Tami, et al., 2007). However, although sachets of pyrethroid insecticides were not only bundled with all bed nets at no extra costs to the recipient, and also made available at subsidized prices through the private retail sector, the insecticide treatment formulations available at the time *Zuia Mbu* which was 2.5% lambda-cyhalothrin SC (capsule suspension) manufactured by Zeneca Ltd UK (Schellenberg et al., 1999), lacked sufficient durability (Erlanger et al., 2004), so measured EIRs remained in triple figures well into the early 2000s (Killeen, Tami, et al., 2007). However, once much longer lasting insecticide treatments became available, initially KO Tab 1-2-3® (Bayer Environmental Sciences at the time, now Envu) (Oxborough et al., 2009; Yates et al., 2005) in 2004 and then Icon MAXX® (Syngenta AG) (Tungu et al., 2015; WHO, 2008) from 2008 onwards, a dramatic reduction in malaria transmission occurred, resulting in an 18-fold reduction of total EIR (Russell et al., 2010) that has been sustained since then (Kaindoa et al., 2017; Lwetoijera et al., 2014; Mapua et al., 2022).

Following the sudden rapid scale up of LLINs, primarily through the sudden improvement in treatment products made available for bed nets, the composition and biting habits of local vector populations shifted dramatically (Russell et al., 2011; Russell et al., 2010). While nominate *An. gambiae* had formerly been the dominant malaria vector in the area, it had already became quite rare by 2008 and then virtually disappeared from mosquito collections in the valley from 2009 onwards (Kaindoa et al., 2017; Lwetoijera et al., 2014; Russell et al., 2011). Despite these gains, malaria has not been completely eliminated, because vector species *An. arabiensis* and *An. funestus s.s.* exhibit different combinations of insecticide resistance traits and evasive behaviours that allow them mediate persisting EIRs varying around a mean of about a dozen infectious bites per person per year (Kaindoa et al., 2017; Lwetoijera et al., 2014).

Although *An. gambiae s.s*. was thought to have been eliminated from the Kilombero floodplain, herein is reported an ecological study of its two more zoophagic sibling species, namely *An. arabiensis* and *An. quadriannulatus* Theobald, spanning an environmentally heterogeneous land use gradient across a previously unsurveyed eastern fringe of the valley (D.R. Kavishe et al., 2025; Walsh et al., 2025) that revealed refuge populations of this species persisting among small frontier communities living at the interface between human settlements and conserved wilderness. These remarkably focal ecological niches where nominate *An. gambiae* still survive appear closely associated with locally common livelihood-driven nocturnal activities, particularly fishing and guarding crops against wild animals throughout the night. However, their apparent absence from both fully conserved natural areas lacking any resident humans, and from well-established villages nearby with more typical human behaviour where most people are reasonably well protected by LLINs, seems to confirm the complete dependence of this species upon safe access to human blood.

## Methods

### Study setting and sampling frame

This repeated rolling cross-sectional study of population ecology within the *An. gambiae* complex was conducted across the Ifakara–Lupiro–Mang’ula (ILUMA) Wildlife Management Area (WMA), some nearby villages immediately to the west, and adjacent areas of Nyerere National Park (NNP) to the east (Figure 1), as previously described in detail elsewhere (D. Kavishe et al., 2025; D.R. Kavishe et al., 2025; Walsh et al., 2025). This ecologically diverse study area was ideally suited to addressing the objectives of several complementary conservation biology (Duggan, 2023; Duggan et al., 2024; L. M. Duggan et al., 2025; Lily M. Duggan et al., 2025), vector ecology (D.R. Kavishe et al., 2025; Walsh et al., 2025) and entomological methodology studies (D. Kavishe et al., 2025). This integrated set of studies collectively aimed to explore whether portfolio effects (Killeen & Reed, 2018) arising from landscape heterogeneities in the availabilities of various potential mammalian blood sources might allow refugia population of malaria vectors to survive in wilderness areas (Walsh et al., 2025), where anthropogenic selection pressure for insecticide resistance may be greatly attenuated. The study area, as described by (D.R. Kavishe et al., 2025; Walsh et al., 2025) spans a geographical gradient of natural ecosystem integrity (Lily M. Duggan et al., 2025) with systematically varying sibling species composition within the *An. gambiae* complex, ranging from fully domesticated land cover with only *An. arabiensis* in the west, through to completely intact natural ecosystems dominated by *An. quadriannulatus* to the east (Walsh et al., 2025).

**Figure 1.**
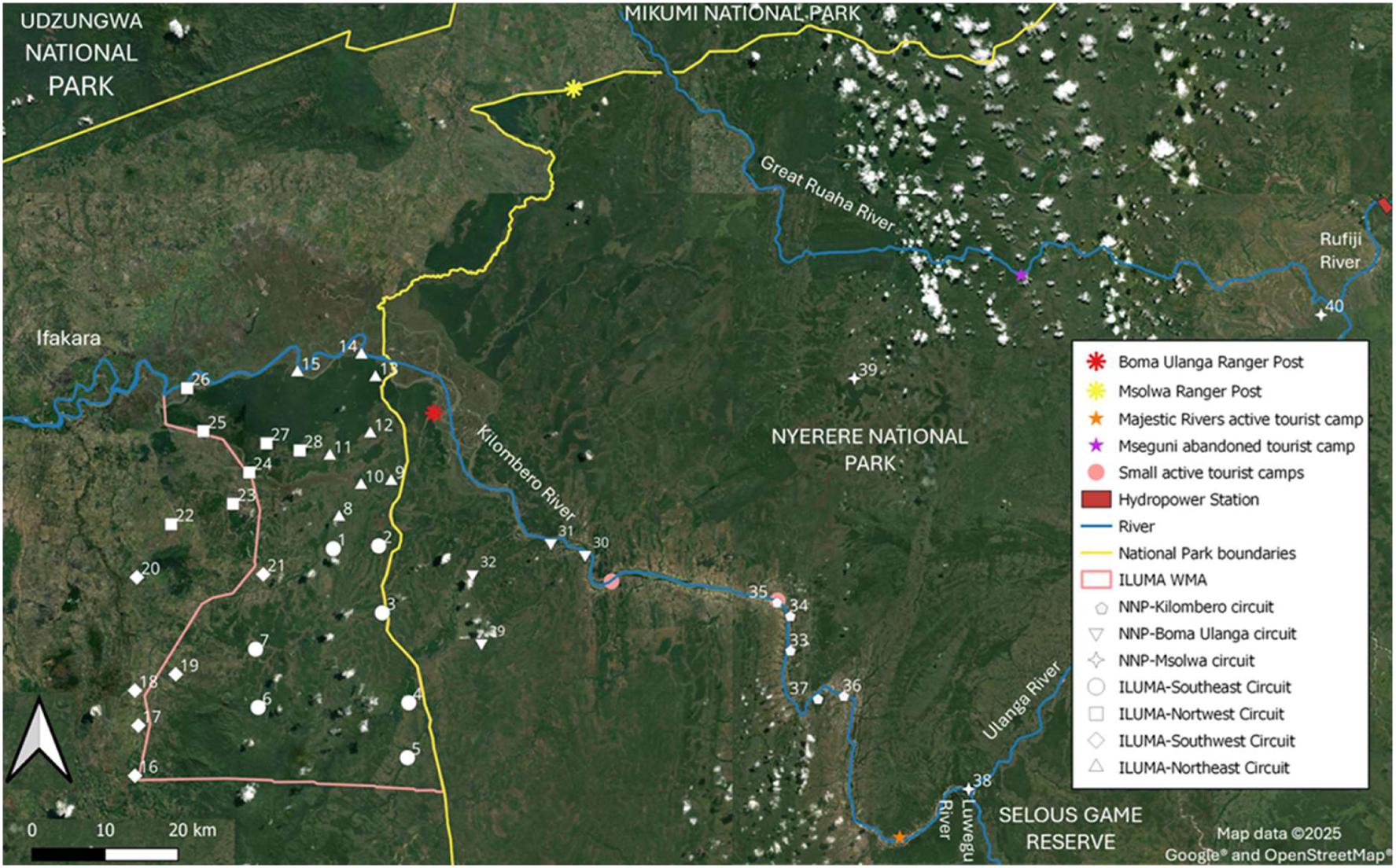
Map displaying the distribution of suitable camping locations used as the sampling frame for all the surveys described herein, illustrated in the geographic context of the boundaries of the Ifakara-Lupiro-Mang’ula Wildlife Management Area (ILUMA WMA), Nyerere National Park (NNP) and the Udzungwa Mountains National Park (D.R. Kavishe et al., 2025; Walsh et al., 2025). For detailed descriptions of the characteristics of each numbered survey location, see previously reported ecological (Duggan, 2023; Duggan et al., 2024; L. M. Duggan et al., 2025; Lily M. Duggan et al., 2025) and entomological studies (D. Kavishe et al., 2025; D.R. Kavishe et al., 2025; Walsh et al., 2025).

Full methodological details, including sampling frame, adult trap deployment and molecular identification protocols have been previously described elsewhere (D. Kavishe et al., 2025; D.R. Kavishe et al., 2025). In brief, adult mosquitoes were repeatedly collected at 40 mobile camps distributed across the ILUMA WMA, nearby villages to the west and adjacent parts of NNP to the east over four survey rounds between January 2022 and December 2023. Adult mosquitoes were captured with either Centre for Disease Control light traps (J.W. Hock, Gainesville Florida, USA) placed at various locations in and around the temporary camps set up by the survey team while visiting each location or with a netting barrier trap (D. Kavishe et al., 2025) set up in a nearby open glade. Adult mosquitoes were identified morphologically as *Anopheles gambiae* sensu lato in the field (Gillies & Coetzee, 1987) and subsequently processed for sibling species identification by polymerase chain reaction (PCR) in the laboratory (Scott et al., 1993; Wilkins et al., 2006).

### Statistical Analysis

Given the strongly anthropophilic nature of *Anopheles gambiae s.s.* (Gillies & Coetzee, 1987; Killeen, Kiware, et al., 2017; White et al., 1972) and its clearly non-random distribution across the study area (Figure 2), the locations surveyed were classified into four categories with obviously distinct ecological human settlement characteristics: (1) Well established villages built legally just outside the conserved area of the WMA, (2) Legally established fishing camps inside ILUMA that have operated since the WMA was established and many years previously, (3) Peripheral scattered settlements inside or immediately outside the western boundary of the WMA, and (4) Locations with largely intact natural land cover and no aggregated settlement of resident humans. These location categories were defined based on observed human settlement patterns, land use and proximity to human activity. For brevity, these categories of locations are respectively referred to hereafter as (1) *Village*s, (2) *Fishing Camps*, (3) *Peripheral settlements* and (4) *Natural Areas* and these classifications are readily related to previous ecological classifications of the area (Duggan et al., 2024; L. M. Duggan et al., 2025; Lily M. Duggan et al., 2025; Walsh et al., 2025).

**Figure 2.**
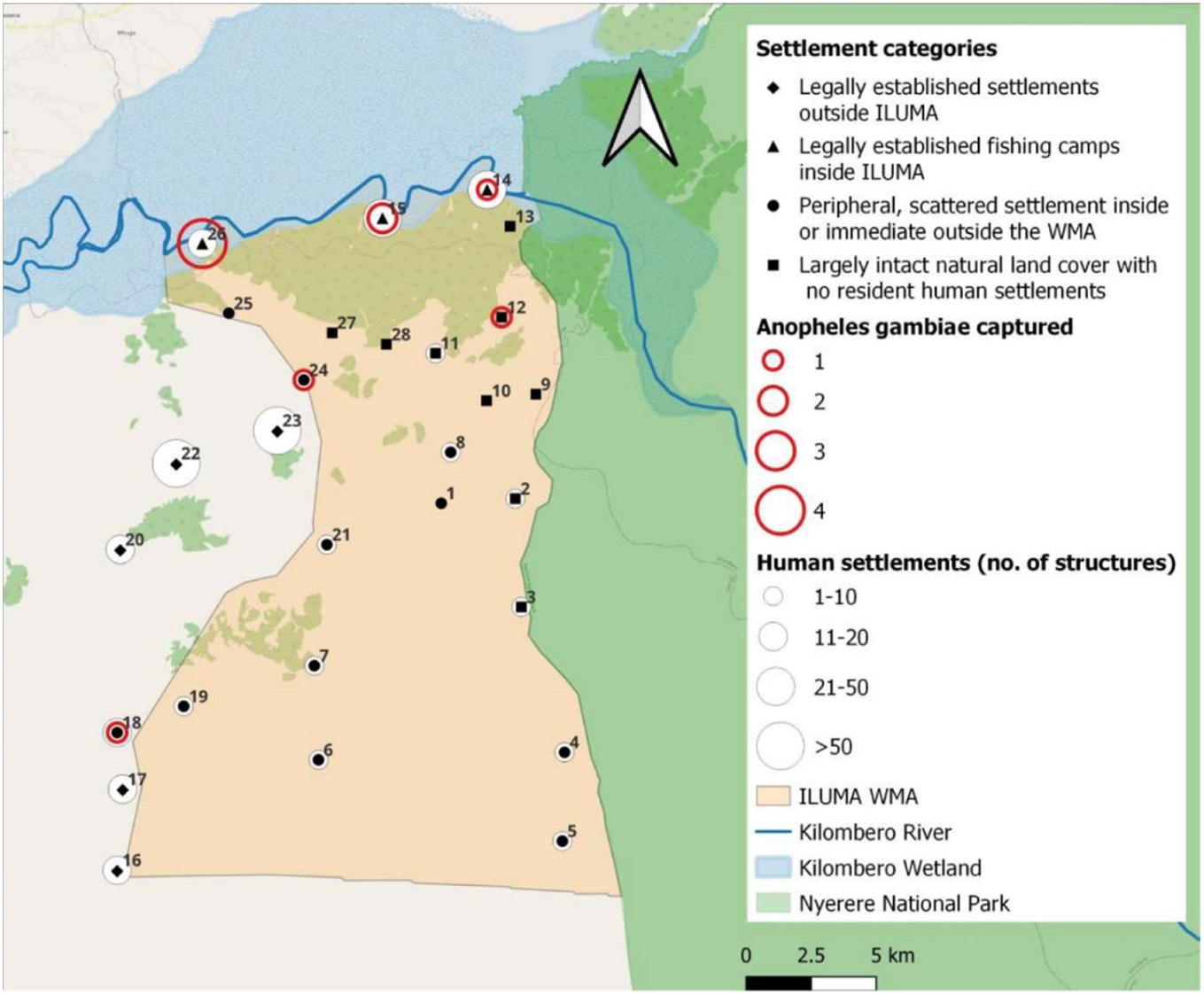
Map displaying the spatial distribution of captured adult *Anopheles gambiae sensu stricto* mosquitoes caught in and around the Ifakara-Lupiro-Mang’ula (ILUMA) Wildlife Management Area (WMA), with each survey location classified into one of four categories with obviously distinct ecological human settlement characteristics: (1) Legally established *villages* with aggregated permanent settlements outside the WMA lands designated for conservation, (2) Legally established *fishing camps* inside ILUMA that have operated since the WMA was established and many years previously, (3) *Peripheral settlements* comprising scattered sets of relatively few houses and/or shelters inside or immediately outside the WMA conservation area, and (4) *Natural areas* with largely intact natural land cover and no aggregated resident settlements inside the WMA or in adjacent parts of Nyerere National Park to the east of it. Note that although 12 additional locations camp 29 to 40 (Figure 1), with fully intact natural land cover and no human settlements inside Nyerere National Park to the east were also surveyed, no *An. gambiae s.s.* mosquitoes were captured there, so this figure focuses specifically on the ILUMA WMA and settled areas immediately to the west of it.

All the following statistical analyses were conducted using *R* version 4.1.3 via *RStudio* version 2023.09.1.494, with version 1.1.9 of the *glmmTMB* package (Team, 2022). To estimate the mean number of *An. gambiae s.s.* mosquitoes caught per trap per night, a generalized linear mixed model (GLMM) without an intercept was fitted to the total count of *An. gambiae s.l.* specimens confirmed to be the nominate species within the complex as the response variable with an assumed zero-inflated Poisson distribution. Settlement category was treated as the sole categorical independent variable, while trapping method (Light trap placed beside occupied tent, light trap within the camp, light trap in a nearby stream, light trap in a nearby open glade or netting barrier trap in a nearby open glade (D.R. Kavishe et al., 2025; Walsh et al., 2025) was included as a random effect. The zero-inflation component was included to allow for the possibility that some locations may have been completely lacking mosquitoes of this sibling species due to prohibitive ecological factors and because it clearly improved the model fit (AIC = 114.84 versus 148.32 for an equivalent model lacking zero inflation). Essentially the same GLMM but with an intercept was also fitted to enable contrasts between different categories of locations, with fishing camps specified as the default reference group. However, neither means nor relative rates could be estimated for the villages due to a complete lack of non-zero observations across all these locations (Figure 2, Table 1), so this category was excluded from the final models presented in table 2.

**Table 1.** Numbers and PCR-confirmed sibling species composition of adult *Anopheles gambiae sensu lato* mosquitoes collected across four settlement categories as illustrated in figure 1: (1) Legally established *villages* with aggregated permanent settlements outside the Ifakara-Lupiro-Mang’ula (ILUMA) Wildlife Management Area (WMA) lands designated for conservation, (2) Legally established *fishing camps* inside the WMA conservation area, (3) *Peripheral settlements* comprising scattered sets of relatively few houses and/or shelters inside or immediately outside the WMA conservation area, and (4) *Natural areas* with largely intact natural land cover and no aggregated resident settlements inside the WMA or in adjacent parts of Nyerere National Park to the east of it.

| Settlement Category | Number of surveyed locations | *An. arabiensis* | *An. gambiae s.s.* | *An. quadriannulatus* | Unknown | Total |
| --- | --- | --- | --- | --- | --- | --- |
| Villages | 5 | 416 | 0 | 5 | 113 | 534 |
| Fishing Camps | 3 | 1790 | 7 | 6 | 484 | 2287 |
| Peripheral Settlements | 11 | 418 | 2 | 4 | 193 | 617 |
| Natural Areas | 21 | 1019 | 1 | 14 | 232 | 1266 |
| Total | 40 | 3643 | 10 | 29 | 1022 | 4704 |

Given that the majority of *An. gambiae s.s.* specimens was captured at the fishing camps inside the WMA conservation area, while none were caught in well-established villages, some further complementary analysis using simpler classification criteria was carried out *post hoc*. The classification of locations was simplified into only two categories to enable inclusion of the data from villages and enhance statistical power: Catch rates at fishing camps versus all locations other than fishing camps were estimated and contrasted using GLMMs that were otherwise identical to those described immediately above.

### Estimation of minimum mean standing population size for *An. gambiae s.s.* around fishing camps

While the scattered peripheral settlements where three *An. gambiae* were caught could not be delineated or enumerated with any confidence, the highly clustered human settlements within the confines of the designated fishing camps represented discrete, well separated and regularly censused human communities that are ideal for estimating the mean sizes of their associated mosquito populations. Borrowing from previous models used to explore mosquito population biodemographic and malaria transmission dynamics (Killeen, 2013; Killeen et al., 2000), the mean standing population size for *An. gambiae s.s.* around each of the three surveyed fishing camps (*N*) was therefore calculated as follows:

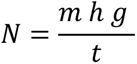

Where *m* is the total number of mosquitoes of that species caught at that location, *h* is the total number of resident humans, *g* is the mean duration of the feeding cycle, which was assumed equivalent to the gonotrophic cycle, and *t* is the total number of trap nights carried out at that location. While the estimates of *h* obtained for Mikeregembe, Mdalangwila and Funga (Camps 14, 15 and 26 and in figures 1 and 2) from the relevant district authorities were approximately 300, 200 and 100, respectively, *t* was recorded as 35, 38 and 29 trap nights, respectively (D.R. Kavishe et al., 2025) and *g* was assumed to be 3 days per gonotrophic cycle (Charlwood et al., 1997). To estimate the sensitivity threshold for the survey procedure, expressed in terms of minimum detectable female population size in the absence of any resident human population competing for the attentions of mosquitoes, *m* was assumed to be 1, *t* to be 40 (maximum possible if all traps operated successfully on all nights during all visits over all four rounds) and *h* to be 10, approximately equivalent to the average size of the survey team.

Note, however, that this calculation assumes that the numbers of mosquitoes caught with each light trap or barrier trap are approximately equivalent to the biting exposure rate of a single human, as is the case for light traps hung beside occupied bed nets (Briët et al., 2015). This may be considered unlikely, even for the light traps hung right beside occupied tents (Service, 1977), and even more so for those placed further away from people or the barrier trap set up away from the camp (D.R. Kavishe et al., 2025; Walsh et al., 2025). Similarly, this calculation assumes that *An. gambiae s.s.* fed only upon humans, which may not necessarily be entirely true, and *g* is likely to be far higher than 3 days now that high bed net coverage forces mosquitoes to undertake longer foraging journeys in search of blood (Birley, 1979; Charlwood & Graves, 1987). These estimates may therefore be considered quite conservative, representing minimum mean standing population sizes, averaged out over the different seasons spanned by the four rounds of surveys.

## Results

A total of 4,704 adult mosquitoes morphologically identified as *Anopheles gambiae s.l.* were collected from all forty surveyed locations, which were each classified as one of four distinct settlement categories: fishing camps, peripheral settlements, natural areas, and villages (Table 1). All specimens were subjected to a polymerase chain reaction (PCR) for sibling species identification, and 78.3% (3682) of them were successfully amplified. Among the successfully amplified adult specimens, 98.9% were *An. arabiensis*, 0.79% were *An. quadriannulatus*, and 0.27% were definitively identified as nominate *An. gambiae sensu stricto*, the primary focus of this analysis (Table 1). All of these PCR-confirmed *An. gambiae s.s.* were captured either within the WMA or immediately adjacent to its western boundary (Figure 2).

The focal concentration of *An. gambiae s.s.* around such a small number of locations in and around the WMA (Figure 2) contrasts starkly with the ubiquitous distribution of its more flexible sibling species (Killeen et al., 2001) *An. arabiensis* across the entire study area (Figure 3). Indeed, *An. arabiensis*, which is thought to feed most readily upon either humans or cattle (Jones, 1980; Killeen et al., 2001; White et al., 1972), were even found in the most isolated locations deep inside NNP that were dozens of kilometres away from the nearest people or livestock (Walsh et al., 2025). Although only a handful of adult *An. quadriannulatus* could be caught (Table 1), because they appear very strictly zoophagic and did not respond to human-baited traps (D. R. Kavishe et al., 2025), larval surveys confirmed that it was ubiquitous across the most intact natural areas in the eastern portion of the WMA and all across NNP (Walsh et al., 2025).

**Figure 3.**
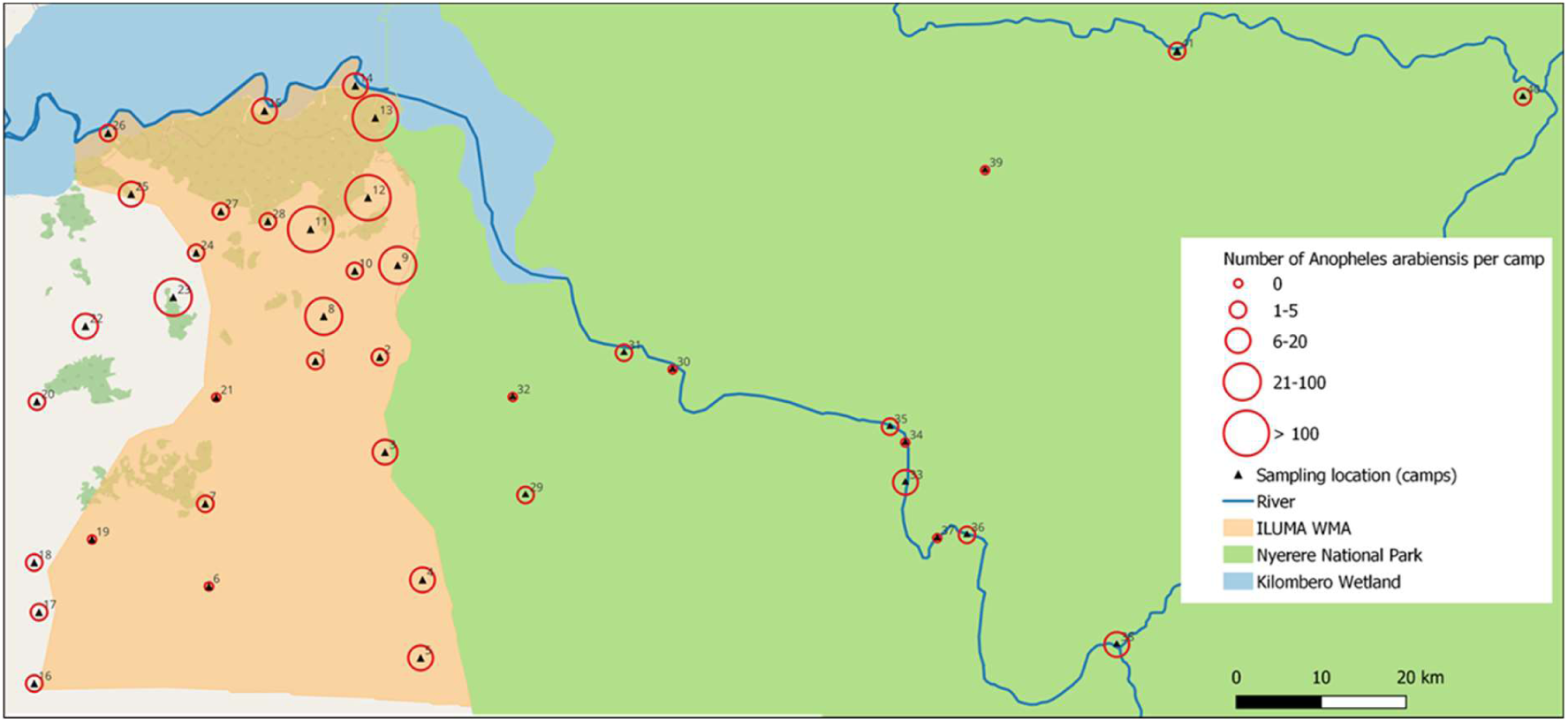
Spatial distribution of adult *Anopheles arabiensis* captured across all survey locations described in Figure 1.

The estimated mean catches of *Anopheles gambiae s.s.* varied significantly across settlement categories, with fishing camps emerging as clear hotspots for this exceptionally efficient vector (Figure 2, Tables 1 and 2). Although only 10 specimens of *An. gambiae s.s.* were caught overall, seven of those were captured in the only three fishing camps inside the WMA that were surveyed. In contrast, this species appeared absent from all 5 well-established villages just outside the WMA and was detected in only two of the 11 peripheral settlements and only one of the 21 ecologically intact natural areas (Figure 1). Fishing camps had substantially higher estimated catches of this species than either of the other two settlement types that could be reliably modelled (Table 2). A *post hoc* GLMM analysis contrasting fishing camps versus all other location types further confirmed the fishing camps as clear hotspots for *An. gambiae s.s.* (Relative Rate [95% Confidence Interval = 18.6 [2.89, 119.0], *P* = 0.002), despite the sparse nature of the data available.

**Table 2.** Estimated mean absolute and relative rates capture of *Anopheles gambiae s.s.* across the four distinct settlement categories, estimated by fitting generalized linear mixed models as described in the *Methods* section

| Location Category | Number of trap nights | Mean catch [95% CI<sup>a</sup>] | Relative Rate [95% CI<sup>a</sup>] | P |
| --- | --- | --- | --- | --- |
| Fishing Camp | 102 | 1.850 [0.397, 8.630] | NA<sup>b</sup> | NA<sup>b</sup> |
| Peripheral settlement | 384 | 0.162 [0.024, 1.090] | 0.088 [0.011, 0.695] | 0.021 |
| Natural areas | 378 | 0.079 [0.007, 0.867] | 0.043 [0.003, 0.495] | 0.012 |
| Village<sup>c</sup> | 172 | NE<sup>c</sup> | NE<sup>c</sup> | NE<sup>c</sup> |

<sup>a</sup> CI: Confidence interval
<sup>b</sup> Not applicable: This was the default reference group for the a priori analysis using a model with an intercept
<sup>c</sup> Not estimable: Data from locations categorized as villages were excluded from the model fitting process because a complete lack of non-zero observations confounded model convergence.

It is interesting that most of the *An. gambiae s.s.* were caught in the three fishing camps, all of which constituted reasonably large, aggregated settlements within the WMA (Figure 4A), it is equally notable none were caught in more typical villages to the west of it (Figure 4B). It is also noteworthy that the other three captured specimens were caught at locations with much sparser, more scattered, far more tentatively established resident human populations (Figure 4C and D) living nearby (Figure 2). Specifically, two were caught near two peripheral settlements on either side of the western boundary of the ILUMA, which both had small resident farming communities that were clearly documented in the surveys (Figure 4C), despite one of these locations (camp 24) being just inside the conservation area of the WMA (Figure 2). While one additional specimen was caught at an ecologically intact natural area much further inside the WMA (camp 12), small resident human settlements were nevertheless documented nearby (Figure 2 and 4D). Specifically, these took the form of a handful of shelters (Figure 4D) hidden within the woodland around the nearest other surveyed location, approximately 3km to the west (Camps 11 in Figures 1 and 2), which is well within the normal flight range of such mosquitoes (Service, 1997).

**Figure 4.**
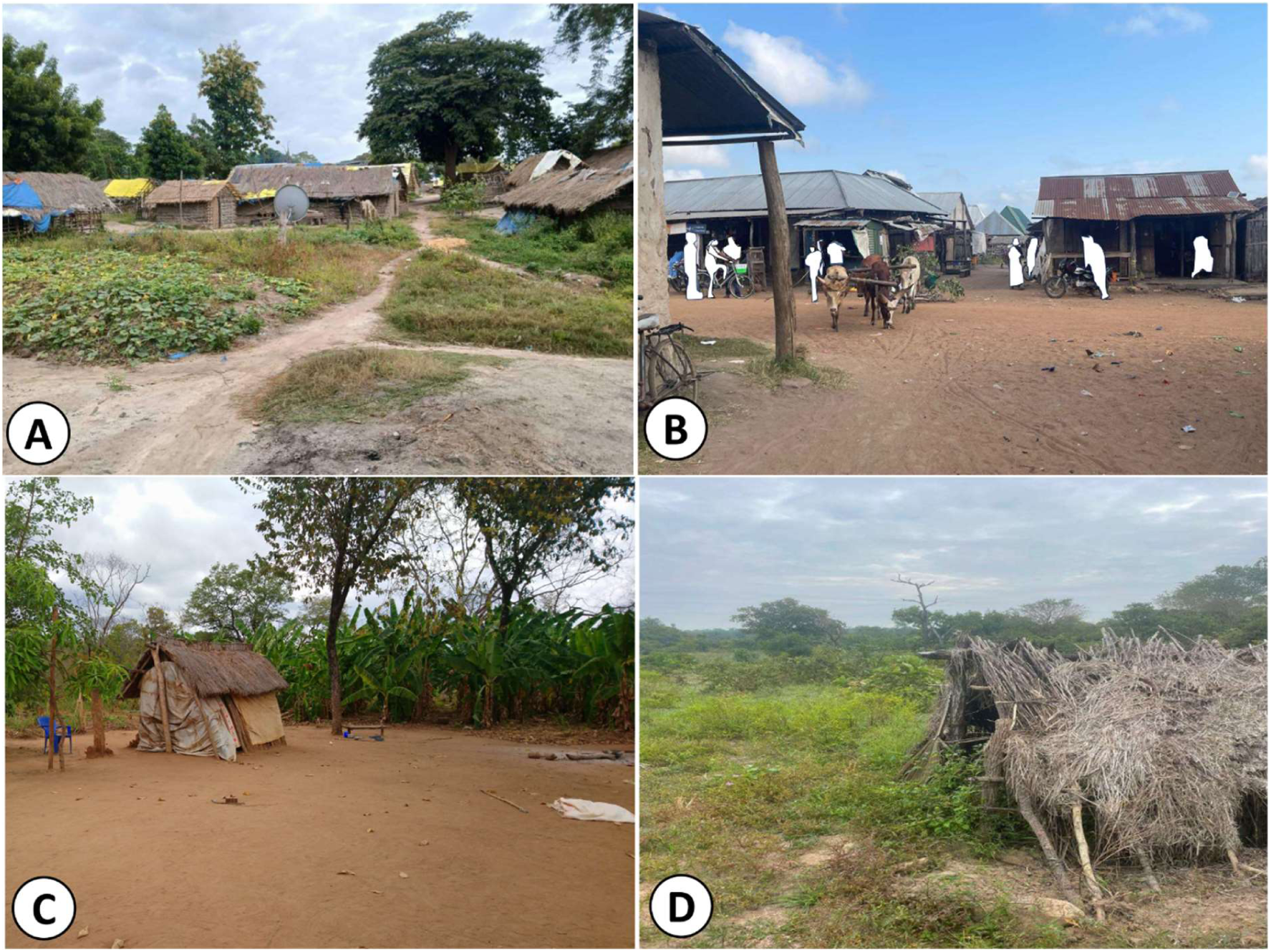
Photographic representations of the categories of locations where *An. gambiae sensu stricto* were and were not detected across the study area, encompassing a land use gradient varying from fully settled and converted lands through to fully intact natural ecosystems (Figures 1 and 2). (A) The fishing camp of Mikeregembe (Location 14 in figures 1 and 2), which is representative of the three surveyed fishing camps where the majority of nominate *Anopheles gambiae s.s.* were captured, all of which represent large, tightly aggregated settlements established inside the Ifakara-Lupiro-Mang’ula (ILUMA) Wildlife Management Area (WMA). (B) A typical well-established village legally built just outside the western boundary of the WMA (Kisaki, Camp number 22 in figures 1 and 2), representative of all five such locations where no *An. gambiae s.s.* were detected. (C) One of the peripheral, sparsely populated farming communities close to the ILUMA boundary (Camp 18 in figures 1 and 2), where one of the two *An. gambiae* s.s. obtained from such locations was caught. (D) The small informal shelters that were found close to camp number 12 (Figures 1 and 2), within three kilometers of the natural woodland site deep inside the WMA where one specimen of *An. gambiae s.s.* was captured.

While it is very plausible that the highly anthropophilic vector species *An. gambiae s.s* may not be able to establish a foothold in wild areas completely lacking stable, resident human populations, it is equally interesting that they could not be found in the best-established villages with the largest human populations (Figures 2 and 3B). What appears to distinguish these fishing camps, peripheral settlements and even one near-natural area with detectable residual populations of *An. gambiae s,s,* from the villages to the west of the WMA is not how many people live there but rather how they live (Figure 5). Specifically, the residents of all these frontier settlements where *An. gambiae s.s.* were caught frequently engage in nocturnal activities that preclude the practical use of insecticidal bed nets (Figure 5). These otherwise unusual livelihood-related human behaviours (Figure 5) were very common among these frontier communities, leaving many residents fully exposed throughout the night and allowing this highly human-specialized malaria vector to feed safely during their preferred peak hours of activity (Killeen, 2014; Killeen et al., 2018; Killeen, Kiware, et al., 2017). Specifically, the fishing activities the authorized camps were established along the Kilombero River to facilitate are largely carried out at night (Figure 5A). Furthermore, farmers living in scattered homesteads along the boundary of the WMA, and often even deep inside it (Duggan et al., 2024; L. M. Duggan et al., 2025; Lily M. Duggan et al., 2025), face the constant threat of crop destruction by wild animals, elephants in particular, so they routinely sleep over their fields in very open shelter structures called *vibanda* (Figure 5B), or even on crude platforms built high up in trees (Figure 5C), where bed net use is physically impractical.

**Figure 5.**
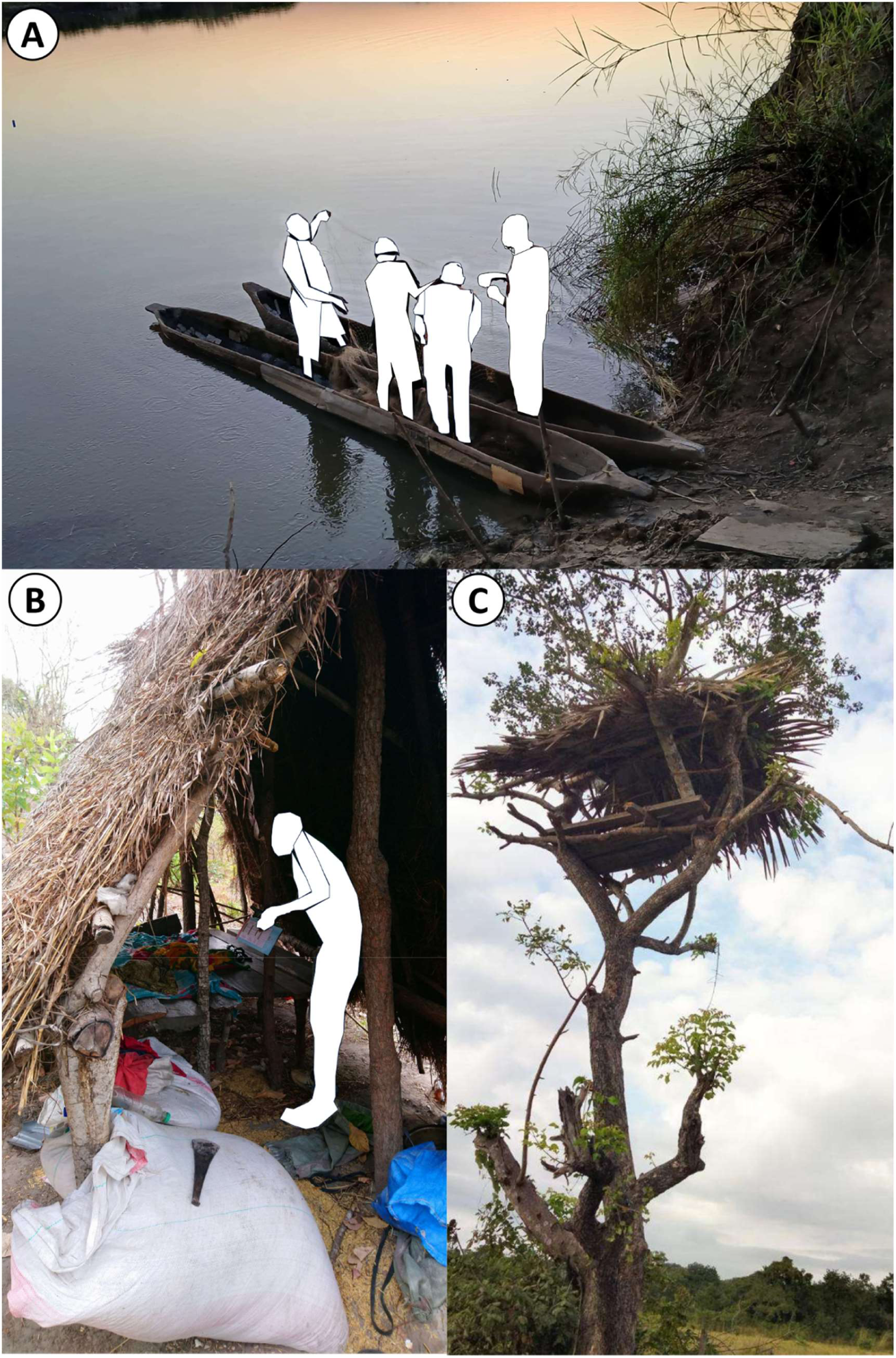
Representative examples of nocturnal livelihood-related activities associated with outdoor exposure to mosquitoes because they preclude the practical use of LLINs, which are commonly practiced among frontier communities living along the eastern fringe of the Kilombero Valley. (A) Fishermen at *Mikeregembe* fishing camp (Location 14 in Figure 1 and 2) preparing at dusk to fish throughout the night along the Kilombero River. (B) Open shelter structures called *vibanda* near the WMA boundary, within which resident farmers sleep overnight to protect their crops from wildlife (Photograph taken near camp 24 in figures 1 and 2). (C) Elevated platforms installed high up in trees, beyond the reach of wild animals, where farmers sleep overnight to guard their crops against them (Photograph taken near camp 24 in figures 1 and 2).

## Discussion

This study provides clear evidence that *Anopheles gambiae sensu stricto*, once considered eliminated from the Kilombero Valley after successful scale-up of LLINs, actually still persists in remarkably focal ecological niches at the interface between human settlement and conserved wilderness. Although the absolute number of captured specimens was small, they nevertheless represent substantial standing populations, estimated as an average of at least several dozen female mosquitoes living at any given time around each of the fishing camps where most of these specimens were caught. Furthermore, in all locations where *An. gambiae s.s.* was detected, residents frequently engaged in unusual nocturnal behaviours related to locally predominant livelihoods, specifically fishing and guarding crops against wildlife, both of which make protective LLIN use difficult or impossible during peak hours of mosquito biting activity.

The clearly non-random spatial distribution of *An. gambiae s.s.* across the study area (Figure 2) strongly indicates that these are quite focal refuge populations, surviving only in specific ecological niches. In these settings, unusually common livelihood-related nocturnal behaviours of their preferred human hosts make LLIN use impractical, thus leaving local residents largely outside the protective reach of conventional vector control tools. Specifically, *An. gambiae s.s.* were only caught at the three surveyed fishing camps established legally within the WMA (Figure 4A), two peripheral farming settlements along its western boundary (Figure 4C) and one well conserved natural location well inside the WMA that nevertheless had a few occupied human dwellings nearby (Figure 4D). While the former were established specifically to enable fishing livelihoods that are largely practiced at night (Figure 5A), the latter three locations where *An. gambiae s.s.* were detected often require local farmers to spend their nights guarding their crops against wild animals, elephants in particular, leaving them fully exposed to mosquitoes in shelters where LLINs are impractical to use (Figure 5B and C).

While the presence of *An. gambiae s.s.* around some of these other frontier settlements (Figure 4C and D) is noteworthy, it is also clear that they were far more readily captured at the fishing camps than anywhere else. This is likely the case because these carefully regulated settlements, legally established with constrained geographic footprints and strictly traditional construction practices within the WMA (L. M. Duggan et al., 2025), represent far more substantial frontier settlements where nocturnal behaviours that are unusually common locally leave residents exceptionally vulnerable to mosquitoes throughout the night. Despite being located within a designated conservation area, all three surveyed fishing camps comprise tight clusters of traditional houses and other open shelter structures because they are, in principle, seasonal settlements, where the construction of permanent structures with durable modern materials is not permitted (Contrast panel A with panel B in figure 4). With every fishing camp having at least 11 established households (Figure 2) all year round, and a resident population that swells far beyond that during peak fishing season, each represents a stable and concentrated aggregation of people who are exceptionally accessible to mosquitoes in an otherwise sparsely populated landscape. As such, they seem to represent ideal refuges for *An. gambiae s.s.*, where it can still access its preferred human hosts in relative safety, despite near-universal coverage with LLINs that has been sustained for almost two decades.

However, the capture of *An. gambiae s.s.* at sites categorized as sparsely populated peripheral settlements (Figure 4C), or even in quite natural areas that nevertheless have a few people living there on a resident basis (Figure 4D), suggests that even very small, scattered human populations are sufficient to support localized persistence of this vector, so long as locally common livelihoods make people a far less hazardous host to attack. In this particular case, residents of such frontier settlements are forced to spend their nights outside the protective reach of the LLINs that most of them would otherwise use (Figure 5B and C). While sleeping in open shelters, such as the *kibanda* illustrated in figure 5B, is widespread across the tropics and has been clearly implicated as a cause of residual malaria transmission in this context (Finda et al., 2019; Killeen, 2014; Monroe et al., 2019; Moshi et al., 2018; Swai et al., 2016) and many others (Edwards et al., 2019; Erhart et al., 2005; Kar et al., 2014; Somboon et al., 1998), spending entire nights on platforms installed high up in trees (Figure 5C) represents an extreme variation on this theme that has not previously been reported to our knowledge.

More importantly, this is the first time to our knowledge that such livelihood-related nocturnal behaviours have not only been implicated as a cause of human exposure to residual malaria transmission (Monroe et al., 2019; Moshi et al., 2018) but also a means for such a highly anthropophagic and exceptionally efficient malaria vector mosquito to persist at sparse but nevertheless self-sustaining population densities in the face of near-universal coverage with the LLINs it is known to be so vulnerable to (Killeen, Kiware, et al., 2017; Killeen et al., 2013; Sinka et al., 2016). In this particular case, it is especially notable because LLINs appear to have eliminated it from most of the rest of the Kilombero Valley (Killeen, Kiware, et al., 2017; Russell et al., 2011), a context in which it was historically the dominant vector (Russell et al., 2010). In relation to strategies for eliminating such exceptionally important anthropophagic malaria vectors, which are as vulnerable to LLINs and IRS as they are efficient as vectors because of their strong preferences for human blood (Killeen, Kiware, et al., 2017; Russell et al., 2010; White et al., 2011), such persisting foci could be appropriately described as *residual vector populations*.

Furthermore, this interpretation appears to be supported by the observation that no *An. Gambiae s.s.* whatsoever were caught at any of the five surveyed farming villages just outside the western boundary of the WMA (Figure 2), all of which have substantial human populations (Figure 4B) practicing more typical livelihoods and construction practices, nor in any intact wilderness areas that lack any resident human populations (Figure 2). Reassuringly, the near complete absence of *An. gambiae s.s.* from intact wilderness areas (Duggan et al., 2024; Lily M. Duggan et al., 2025) deep inside the WMA (Figure 2) or the national park to the east of it (Figure 1) seems to further confirm the absolute dependence of this highly specialized species upon humans as their sole source of blood. Indeed, the only exception appears to prove the rule: While no signs of human settlement were observed immediately around camp 12, where the only *An. gambiae s.s.* specimen was caught in any intact natural area, a few unauthorized homesteads with minimal shelter (Figure 4D) were found in the vicinity of camp 11, approximately 3km to the west (Figure 2) and well within the routine flight range of host-seeking mosquitoes (Service, 1997). In stark contrast, this same set of surveys found that *An. arabiensis* (Figure 3) and *An. quadriannulatus*, both of which are known to feed readily upon animals, were ubiquitous across the wildest parts of the study area, even in places deep inside NNP that were dozens of kilometres away from the nearest humans or livestock (Walsh et al., 2025).

Furthermore, one of the more striking results reported herein is the apparent absence of *An. gambiae s.s.* from the five formally recognized villages, despite these areas having larger human populations than the fishing camps with far more secure tenure of residence. This apparent paradox suggests that vector control efforts are particularly effective against this mosquito species in conventional rural settings where human behaviours align reasonably well with the assumptions of indoor-focused interventions (Bhatt et al., 2015; Killeen et al., 2006; West et al., 2014). However, like almost all other villages across the tropics, alignment with these assumptions is imperfect, with mosquitoes often accessing human blood despite high LLIN usage, through a diversity of routine household or social activities in the evenings and early mornings, not to mention varying minor fractions of people who sleep without a net, sometimes outdoors (Finda et al., 2019; Monroe et al., 2019; Moshi et al., 2018). In these more typical, well-established villages, it seems that the benefits of numerous feeding opportunities created by all the usual human behaviours (Erhart et al., 2005; Hii et al., 2021; Kar et al., 2014; Somboon et al., 1998) that ubiquitously facilitate residual malaria transmission (Durnez & Coosemans, 2013) are outweighed by the high risks of being killed when they do encounter people sleeping under protective LLINs. In this particular geographic context, even the sparsest of human populations appears sufficient to enable focal survival of *An. gambiae s.s.*, so long as the mass effects of LLINs on vector populations are attenuated by locally common nocturnal activities that undermine their practical utility (Figure 5).

Overall, it seems that *An. gambiae s.s.* not only needs humans to be present but also that it can safely access them. A natural corollary of this conclusion is that this vector species is an exclusive obligate ectoparasite of human beings that should be possible to eliminate more widely if gaps in the effective coverage achieved by existing insecticidal vector control measures, namely LLINs and IRS, can be closed with complementary new interventions like transfluthrin emanators (Achee et al., 2012; Nakyaze et al., 2024; Vajda et al., 2024) and endectocide treatments (Chaccour et al., 2025).

However, a more worrisome implication of these persisting residual populations of *An. gambiae s.s.* is the evolutionary potential for emergence of robust insecticide resistance traits and behavioural adaptations that undermine hard-earned gains to date (Ranson & Lissenden, 2016; WHO, 2012). The virtual disappearance of this sibling species from most villages in this part of south-eastern Tanzania is attributed to successful LLIN distribution progammes, which decisively shifted the composition of the local malaria vector guild toward the more zoophagic, outdoor-biting, cautious species *An. arabiensis* (Kitau et al., 2012; Russell et al., 2011). However, the survival of *An. gambiae s.s.* at such low but nevertheless self-sustaining population densities, in foci scattered along fringes of human existence, indicates that this highly anthropophagic species has found partial refuge from the hazards of widespread LLIN use, creating quite dangerous evolutionary pressures (Hancock et al., 2020; Hastings et al., 2022; Ranson & Lissenden, 2016; WHO, 2012). Extensive insecticide use in public health has imposed strong selection pressure on African malaria vectors (Ranson & Lissenden, 2016; Ranson et al., 2011), leading to widespread pyrethroid resistance and changes in biting habits across many regions (Bayoh et al., 2014; Moiroux et al., 2012; Russell et al., 2011). In Tanzania, for example, resistance to all four major classes of insecticides used in nets and sprays had already been documented within the *An. gambiae* complex a decade ago (Kisinza et al., 2017). Such residual populations of *An. gambiae s.s.* that persist despite high net coverage could act as reservoirs from which this notorious vector could rebound if control efforts were to slacken (Cohen et al., 2022), or more troublesome physiological (Ranson et al., 2011; WHO, 2012) and/or behavioural resistance (Govella et al., 2013) traits were to emerge because such dangerous partial intervention pressure creates optimal conditions for their evolutionary selection (Hastings et al., 2022; Hemingway & Ranson, 2000). In short, the presence of even a handful of *An. gambiae s.s.* among frontier communities living in and adjacent to conserved wilderness areas is not only of immediate epidemiological significance there and now but could also prove to be of exceptional strategic importance over the long term. There may therefore be much to be gained by boldly tackling these last-mile challenges in such marginalized human settlements, where refuge populations of some of the world’s most dangerous, human-dependent malaria vectors still persist outside of the traditional indoor, village-focused paradigm (Govella & Ferguson, 2012) but could well be nudged into local extinction by appropriately targeted new interventions (Killeen, 2014; Killeen et al., 2013).

Having said that, several substantial limitations of this study must be acknowledged. The very low absolute number of *An. gambiae s.s.* caught limited the certainty of the conclusions reached, even though appropriate statistical tests indicate several of them appear probable. For example, the identification of fishing camps as hot spots for *An. gambiae s.s.* persistence relied on only seven mosquitoes collected from only three surveyed camps, and only one other specimen was obtained from each of three other collection sites. While such a sparse dataset obviously limited statistical power and certainty of interpretation, it is nevertheless encouraging that the geographic distribution of this now-rare vector species was so clearly non-random and that some reasonably clear specific characteristics of such refuge populations were identified that could be acted upon.

However, regarding the apparent absence of *An. gambiae s.s.* from most locations across the study area, absolute sampling sensitivity of the survey procedure merits consideration. With a maximum of only eight nights of sampling at each camp location, and most of the traps placed too far away from people to attract human specialized mosquitoes as efficiently as they otherwise could be (D.R. Kavishe et al., 2025; Walsh et al., 2025), mosquito populations big enough to sustain themselves could have evaded detection at any of these individual locations. While the estimated minimum detectable standing population size for these surveys was slightly less than one female for places without any hosts other than the survey team, that estimated sensitivity threshold increases approximately in proportion to the size of local resident human population competing for the attentions of mosquitoes (Eq. 1). It is therefore possible that modestly sized self-sustaining populations of *An. gambiae s.s.* around some of these surveyed locations, especially the largest villages, were missed that would require more intensive sampling and/or sensitive trapping methods to reliably detect.

Having said that, these findings highlight a nice example of how much can be learned from creative adaptation of standard entomological surveillance approaches for African malaria vectors, which tend to focus on indoor collections centred around relatively accessible human settlements. Such sampling frameworks may underestimate or entirely miss epidemiologically relevant vector populations that persist at low densities in frontier communities living at the fringes of human interaction with conserved natural ecosystems, where they may also act as bridge vectors for a diversity of zoonoses (de Alvarenga et al., 2015; Killeen, 2014; Killeen, Tatarsky, et al., 2017).

### Conclusions

In conclusion, our findings show that small, highly focal populations of *Anopheles gambiae* s.s. persist on the fringes of the Kilombero Valley in ecological niches where human behaviour leaves people exposed to mosquito bites at night. These residual vector populations were found almost exclusively in fishing camps and scattered frontier settlements inside or adjacent to wildlife conservation areas, where livelihood activities such as night fishing and crop guarding limit effective use of bed nets. Consequently, this strictly human-dependent mosquito continues to survive in these ecological refuges, despite being virtually eliminated from neighbouring villages with long-term use of LLINs. These observations highlight the urgent need for complementary vector control tools that extend protection beyond indoor settings. Spatial repellents and other area-wide outdoor interventions offer promising options to reduce exposure in such open environments (Achee et al., 2012; Nakyaze et al., 2024; Vajda et al., 2024). Addressing these last pockets of *An. gambiae* s.s. is not only vital for protecting underserved communities, but also for preventing these mosquitoes from acting as reservoirs for malaria resurgence or focal points for the evolution of insecticide resistance under sustained partial intervention pressure (Ranson & Lissenden, 2016). Critically, future surveillance efforts must adopt a more flexible and ecologically sensitive design capable of detecting and responding to these residual vector niches effectively.

## AUTHOR CONTRIBUTIONS

Deogratius R. Kavishe: Conceptualization, investigation, methodology, formal analysis, writing original draft of the manuscript. Lucia J. Tarimo: Implementation, maps preparations, review, editing, and validation. Rogath V. Msoffe: Supervise molecular identification, Review, editing, and validation. Katrina A. Walsh, Lily M. Duggan and Fidelma Butler: Review, editing, and validation. Emmanuel W. Kaindoa: Review, editing, and validation. Gerry F. Killeen: Acquisition of fund, Conceptualization, investigation, visualization, methodology, formal analysis, review, editing and validation. All authors read and approved the final submitted version of the manuscript.

## ACKNOWLEDGEMENTS

The authors wish to thank the Village Game Scouts of ILUMA WMA, for their hard work and participation in field activities. We also thank all the staff, management and stakeholder communities of the ILUMA WMA for all their collaboration and kind assistance over the course of the study. Furthermore, we thank Mr Frederic Masanja, Prof Honorati Masanja, Mr Fadhili Sango, Ms Elaine Kelly, Dr Ronan Hennessy, Ms Leonie O’Doherty, Ms Sonia Montero and Prof Sarah Culloty for all the essential institutional support provided by the Ifakara Health Institute and University College Cork over the course of the study. A very special word of thanks is due to our deceased friend and colleague, Mr Octavian Malopola, without whom this work could never have begun, much less completed safely and successfully.

## ETHICAL CONSIDERATIONS

Ethics approvals for this study were obtained from the Institutional Review Board of the Ifakara Health Institute (Ref no. IHI/IRB/No: 5-2021) and the Medical Research Coordinating Committee (MRCC) at the National Institute of Medical Research in Tanzania (Ref. NIMR/HQ/R.8a/Vol. IX/3719). Because the field sites were inside the wildlife management area and national park the research also was approved by Tanzania Wildlife Research Institute (TAWIRI-Ref. No. AB.235/325/01/37), Tanzania Wildlife Management Authority (TAWA-Ref. No. AE.542/712/01), and Tanzania National Parks (TANAPA-BE.161/376/01).

## CONSENT FOR PUBLICATION

Permission to publish was granted by The National Institute of Medical Research (NIMR), Tanzania Ref no. BD.242/437/01C/101

